# Forecasting dengue epidemics using a hybrid methodology

**DOI:** 10.1101/498394

**Authors:** Tanujit Chakraborty, Swarup Chattopadhyay, Indrajit Ghosh

## Abstract

Dengue case management is an alarmingly important global health issue. The effective allocation of resources is often difficult due to external and internal factors imposing nonlinear fluctuations in the prevalence of dengue fever. We aimed to construct an early-warning system that could accurately forecast subsequent dengue cases in three dengue endemic regions, namely San Juan, Iquitos, and the Philippines. The problem is solely regarded as a time series forecasting problem ignoring the known epidemiology of dengue fever as well as the other meteorological variables. Autoregressive integrated moving average (ARIMA) model is a popular classical time series model for linear data structures whereas with the advent of neural networks, nonlinear structures in the dataset can be handled. In this paper, we propose a novel hybrid model combining ARIMA and neural network autoregressive (NNAR) model to capture both linearity and nonlinearity in the datasets. The ARIMA model filters out linear tendencies in the data and passes on the residual values to the NNAR model. The proposed hybrid approach is applied to three dengue time-series data sets and is found to give better forecasting accuracy in comparison to the state-of-the-art. The results of this study indicate that dengue cases can be accurately forecasted over a sufficient time period using the proposed hybrid methodology.

## 1. Introduction

Dengue incidence has increased drastically in recent decades, putting almost half of the humans at risk (1). Dengue fever (DF) is caused by four closely related serotypes (DEN-1, DEN-2, DEN-3, and DEN-4) of dengue virus. The virus is transmitted to humans by the bites of infected female Aedes Aegypti and Aedes Albopictus mosquitoes (2). Although the case fatality rate of DF is meager, it can progress to severe complications such as dengue hemorrhagic fever (DHF) and dengue shock syndrome (DSS).

The dengue endemic countries of Asia and Latin America have reported that DHF and DSS are among the leading causes of hospitalization and death therein. The prevalence of DF is geographically dispersed over the countries situated in the tropical and subtropical regions (2; 3), with local variations in risk and circulation of multiple dengue serotypes. However, no medicine is still available to cure DF; treatment only includes medications for clinical symptoms. Although some candidate vaccines (including live attenuated mono, tetravalent formulation inactivated whole virus vaccines, and recombinant subunit vaccines) are undergoing various phases of clinical trials but none of the vaccines are yet employed (4; 5).

Consequently, health care officials have to rely on early warning systems in order to optimally disseminate available resources in the dengue-prone areas. Impact of severe complications of DF can be reduced by effective case management in endemic regions (6). Policy-makers can employ preventive measures and allocate sufficient resources in the areas with the highest epidemic risk, whenever one can notify the local health care officials on time. Thus, forecasting the future dengue incidence is of utmost interest. Time series forecasting tools are suited when very little knowledge is available regarding the data generating process. Previously, various attempts were made to predict dengue epidemics with several degrees of success (see (6; 7; 8; 9; 10; 11; 12)). These attempts to anticipate the future dengue cases have increased our understanding and have given some insights into further challenges in forecasting epidemics.

Among various traditional models, ARIMA is popularly used for forecasting linear time series. Machine learning models such as artificial neural Networks (ANN) and support vector machines (SVM) are proved to perform well for nonlinear data structures in time series data. Dengue datasets are neither purely linear nor nonlinear. They usually contain both linear and nonlinear patterns. If this is the case, then the individual ARIMA or ANN are inadequate to model such situations. Therefore the combination of linear and nonlinear models can be well suited for accurately modeling such complex autocorrelation structures. Several hybrid methodologies were discussed in previous literature to solve a variety of time series problems arose in econometrics, financial stocks, electricity and other applied areas (13; 14; 15; 16; 17; 18; 19; 20; 21). Zhang’s hybrid ARIMA-ANN model (22) has gained popularity due to its capacity to forecast complex time series accurately. The pitfall of hybrid ARIMA-ANN model lies in the selection of the number of hidden layers in the ANN architecture that involves subjective judgment. To ignore this drawback we took recourse to the NNAR model which is more of a “white-box-like” approach. NNAR fits a feed-forward neural network model with only one hidden layer to a time series with lagged values of the series as inputs (also flexible to handle some other exogenous data). The advantage of NNAR over ANN is that NNAR is a nonlinear autoregressive model, and it provides less complexity, easy interpretability and better prediction as compared to ANN. Due to this NNAR is getting more attention in recent literature of non-stationary time series forecasting (23).

The primary motivation of this paper is to handle the time series data with several decisions regarding how we describe the recent dynamics of the observed values of the series. Taking a final decision in policy making based on a component model may be dangerous in such a severe problem like dengue forecasting where one frequently observe changes in the dynamic properties of the variable being measured. Hybridization of two or more models are most common solution to this problem where one can take advantages of diversity among models to reduce both the bias and variances of the prediction error obtained using single models (24). Even from the practitioners point of view, hybrid models are more effective when the complete data characteristics are not known (25). Motivated from these discussions, this paper proposes a novel hybrid ARIMA-NNAR model that captures complex data structures and linear plus nonlinear behavior of dengue datasets. In the first phase of our proposed model, ARIMA catches the linear patterns of the dataset. Then the NNAR model is employed to capture the nonlinear patterns in the data using residual values obtained from the base ARIMA model. The proposed model has easy interpretability, robust predictability and can adapt seasonality indices as well. Through experimental evaluation, we have shown the excellent performance of the proposed hybrid model for the dengue epidemics forecasting for three different regional datasets.

## 2. Methodology

The deficiencies of the single time series models can be overcome with the hybrid methodology. Achieving stationarity in both the mean and variance is considered essential in classical time series forecasting methods but the literature of machine learning methods are capable of effectively modeling any type of data patterns and can therefore be applied to the original data (26). In the process of capturing typical patterns in the data, a combination of linear and non-linear time series model is often applied to showcase the salient features of the data sets. Few popular hybrid models in this literature are hybrid ARIMA-ANN (22; 18) and hybrid ARIMA-SVM model (19) where the main motivation was to understand both linear and nonlinear patterns of the time series data. These models have shown better performance in terms of prediction accuracy for different forecasting problems of economics, sales, finance, carbon price, stock, electricity, etc. In this work, we propose a novel hybridization of ARIMA and NNAR model to solve the dengue forecasting problem. Below we give a brief description of the component models used in the hybridization.

### 2.1. ARIMA Model

The ARIMA model, introduced by Box and Jenkin (27), is a linear regression model indulged to track linear tendencies in stationary time series data. The model is expressed as ARIMA(p,d, q) where p, d, and q are integer parameter values that decide the structure of the model. More precisely, p and q are the order of the AR model and the MA model respectively, and parameter d is the level of differencing applied to the data. The mathematical expression of the ARIMA model is as follows

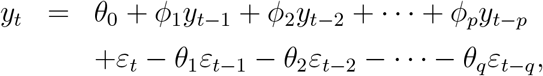

where *y_t_* is the actual value, *ε_t_* is the random error at time *t*, *ϕ_i_* and *θ_j_* are the coefficients of the model. It is assumed that *ε_t−l_*(*ε_t−l_* = *y_t−l_* – *ŷ_t−l_*) has zero mean with constant variance, and satisfies the i.i.d condition. The methodology consists of three iterative steps: (1) model identification and model selection; (2) parameter estimation of the model parameters, (3) model diagnostics checking (namely, residual analysis) are performed to find the ‘best’ fitted model.

In the model identification and model selection step, differencing is applied once or twice to achieve stationarity for non-stationary data. As stationarity condition is satisfied, the autocorrelation function (ACF) plot and the partial autocorrelation function (PACF) plot are examined to select the AR and MA model types. The parameter estimation step involves an optimization process utilizing metrics such as the Akaike Information Criterion (AIC) and/or the Bayesian Information Criterion (BIC). Finally, in the model checking step, the residual analysis is carried out to finalize the ‘best’ fitted ARIMA model. ARIMA model is a data-dependent approach that can adapt to the structure of the data set. But it has the major disadvantage that any significant nonlinear data set can restrict the ARIMA model. Therefore, the proposed hybrid model uses NNAR model to deal with the nonlinear data patterns for forecasting complex time series structure.

### 2.2. NNAR model

Neural nets are based on simple mathematical models of the brain, used for complex nonlinear forecasting. A neural network can be thought of as a network of “neurons” that are arranged in layers (viz. input, hidden and output layers). The forecasts are obtained by a linear combination of the inputs. The weights are selected in the network model using a “learning algorithm” that minimizes the mean squared error.

NNAR model is a nonlinear time series model which uses lagged values of the time series as inputs to the neural network (23). NNAR(p,k) is a feed-forward neural networks having one hidden layer with p lagged inputs and k nodes in the hidden layer. For example, an NNAR(9,5) model is a neural network that uses the last nine observations (*y*_*t*−1_, *y*_*t*−2_, …, *y*_*t*−9_) as inputs for forecasting the output with five neurons in the hidden layer. This model can be applied to the original nonlinear data without putting any restrictions on the parameters to ensure stationarity. An NNAR(p,k) model uses p as the optimal number of lags (calculated based on the AIC value) for an AR(p) model and k is set to 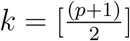 for non-seasonal data sets. To forecast the time series, the NNAR model is applied iteratively with the logistic activation function within the network. For one step ahead forecast, we can utilize the available historical inputs whereas for two steps ahead forecast, we need to use the one step ahead forecast as an input, along with the historical data. This process proceeds until all the required forecasts are computed.

### 2.3. Formulation of the hybrid model

ARIMA model is one of the traditional statistical models for linear time series prediction. On the other hand, the NNAR model can capture nonlinear trends in the data set. So, the two models are consecutively combined to encompass both linear and nonlinear tendencies in the model (28). A hybrid strategy that has both linear and nonlinear modeling abilities is a good alternative for forecasting dengue cases. Both the ARIMA and the NNAR models have different capabilities to capture data characteristics in linear or nonlinear domains. Thus, the hybrid approach can model linear and nonlinear patterns with improved overall forecasting performance. There exist numerous time series models in the literature, and several research shows forecast accuracy improves in hybrid models. The aim of developing a novel hybridization is to harness the advantages of single models and reduce the risk of failures of single models. The underlying assumption of the hybrid approach based on linear and nonlinear model assumption is that the relationship between linear and nonlinear components are additive. The strength of single models for hybridization is very important, and this selection is essential to show the consistent improvement over single models. This paper presents a novel hybridization of ARIMA and NNAR to overcome the limitation of single models and utilize their strengths. With the combination of linear and nonlinear models, the proposed methodology can guarantee better performance as compared to the component models.

The hybrid model (*Z_t_*) can be represented as follows

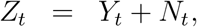

where *Y_t_* is the linear part and *N_t_* is the nonlinear part of the hybrid model. Both *Y_t_* and *N_t_* are estimated from the data set. Let, *Ŷ_t_* be the forecast value of the ARIMA model at time t and *ε_t_* represent the residual at time t as obtained from the ARIMA model; then

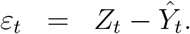

The residuals are modeled by the NNAR model and can be represented as follows

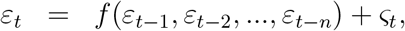

where *f* is a nonlinear function modeled by the NNAR approach and ϛ*_t_* is the random error. Therefore, the combined forecast is

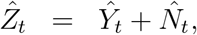

where, 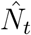 is the forecast value of the NNAR model. The rationale behind the use of residuals in the diagnosis of the sufficiency of the proposed hybrid model is that there is still autocorrelation left in the residuals which ARIMA could not model. This work is performed by the NNAR model which can capture the nonlinear autocorrelation relationship.

In summary, the proposed hybrid ARIMA-NNAR model works in two phases. In the first phase, an ARIMA model is applied to analyze the linear part of the model. In the next stage, an NNAR model is employed to model the residuals of the ARIMA model. The hybrid model also reduces the model uncertainty which occurs in inferential statistics and forecasting time series. A flowchart of the hybrid ARIMA-NNAR model is presented in Figure 1.

**Figure 1:**
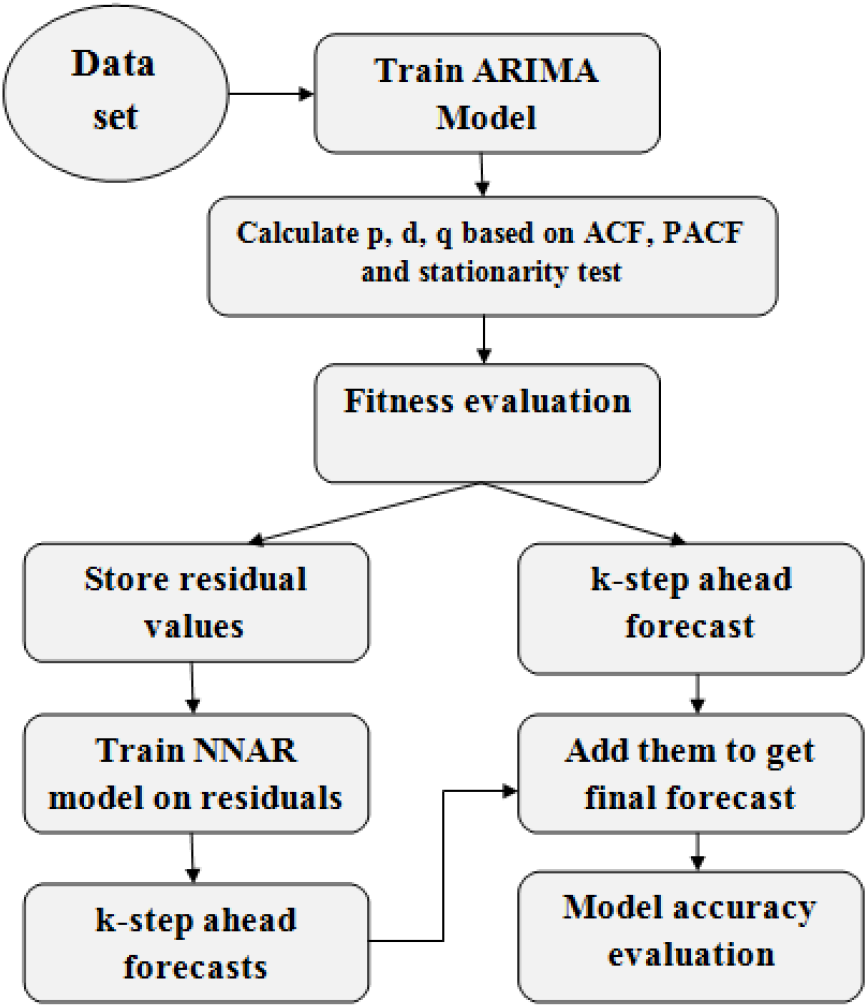
Flow diagram of the proposed model

## 3. Experimental Evaluations

In this section, three popular open-access dengue data sets, namely San Juan, Iquitos and the Philippines data are used to determine the effectiveness of the proposed model. The properties of these data sets are different and have been used in previous studies (7; 10). Different linear and nonlinear models have been studied on these data sets that shows highly nonlinear patterns in these regions. Mean absolute error (MAE); root mean square error (RMSE) and symmetric Mean Absolute Percent Error (SMAPE) are used to evaluate the performances of the proposed model and other single models for dengue data sets.

### 3.1. Data sets

Among the three datasets, two are weekly dengue incidence data, and one is monthly data. For the endemic regions San Juan and Iquitos, weekly laboratory confirmed cases for the time periods from May 1990 through October 2011 and from July 2000 through December 2011, respectively are considered in this study. Weekly dengue incidence data for the San Juan region and Iquitos region are made available in this link http://dengueforecasting.noaa.gov/. The Philippines dataset contains the monthly recorded cases of dengue per 100,000 population in the Philippines. Monthly incidence of dengue in the Philippines is collected from kaggle (see the following link below: https://www.kaggle.com/grosvenpaul/dengue-cases-in-the-philippines/) for the time period January 2008 through December 2016. The Philippines monthly dataset contains a total of 108 monthly observations and we use the total cases reported from all regions in the Philippines in this study. San juan weekly dataset contains a total of 1144 observations whereas Iquitos dataset contains only 520 observations.

### 3.2. Performance measures

The metrics used in this study to evaluate the performance of different forecasting models (including the proposed model) are RMSE, MAE and SMAPE (29).

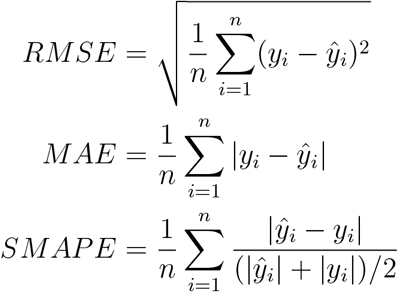

where *y_i_* is the target output, *ŷ_i_* is the prediction and *n* denotes the number of data points. By convention, the lower the value of these metrics, the better the forecast model is.

### 3.3. Analysis of results

We have divided three dengue datasets into training and testing data. For weekly datasets we have kept last six months data for testing model accuracy. We have kept one year data as test data for the Philippines dataset. The behavior of the dataset can be regarded as nonlinear and non-gaussian. The time series plot of the data set show a cyclical pattern with a mean cycle of about 1 year (see Table 1). We have studied ARIMA, ANN, NNAR model for this data. The data set is divided into two samples of training and testing to assess the forecasting performance of the proposed model. For example, the training set of Philippines contains 96 observations (January 2008 – December 2015), is exclusively used for model building. Further, the last 12 month’s data (January 2016 – December 2016) are used for model evaluation. We have applied our proposed hybrid ARIMA-NNAR model to all the three datasets as follows.

**Table 1:**
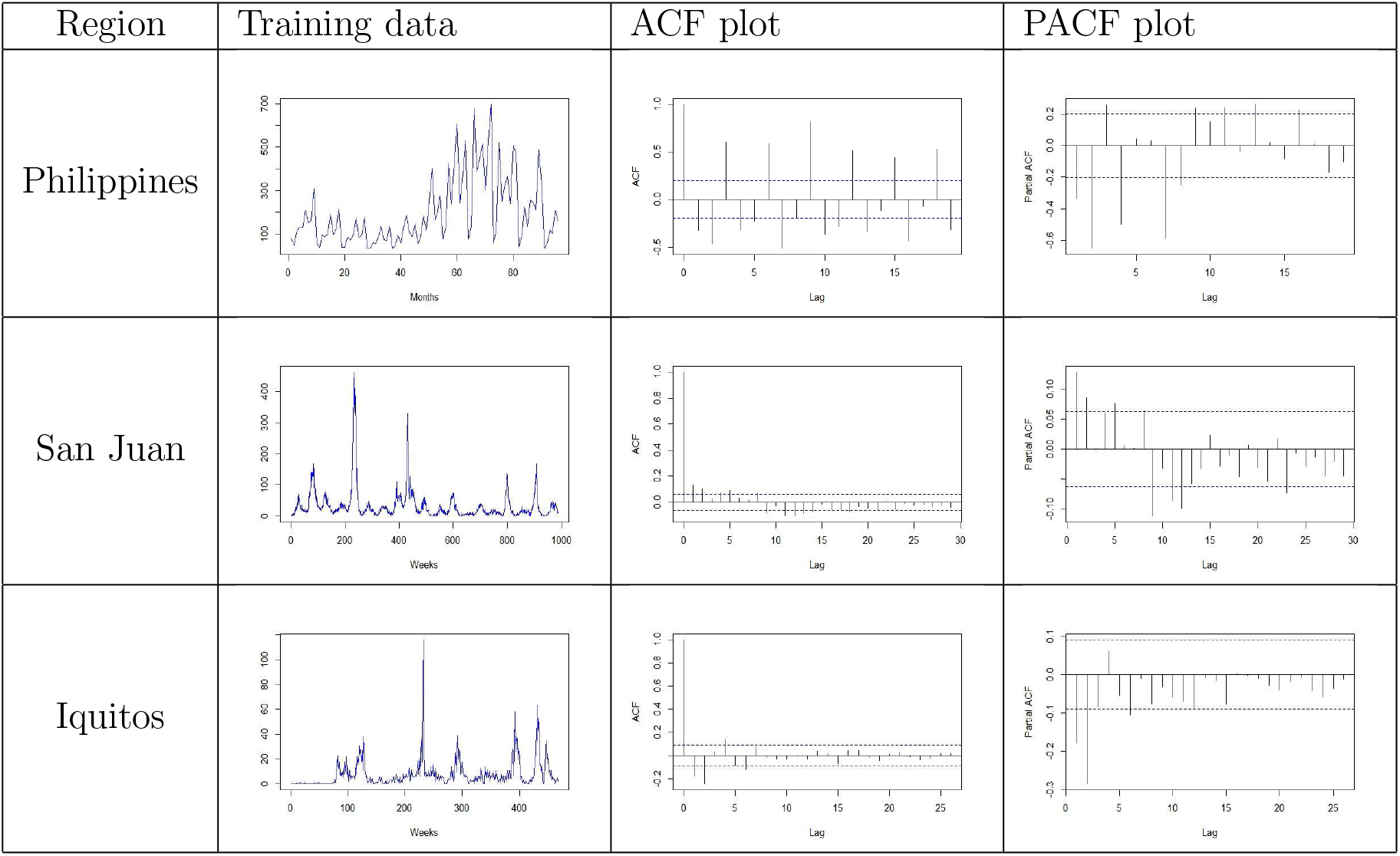
ACF and PACF plots

Linear modeling is done with ARIMA(p,d,q) using “forecast” package in R statistical software. Nonlinear modeling with NNAR approach is done with “caret” package using “nnetar” function in R statistical software. Before fitting an ARIMA model, the order of the model must be specified. The ACF plot and the PACF plot aid the decision process. We choose the ‘best’ fitted ARIMA model using AIC value, for each train data set. The method we use to compute the log-likelihood function for the AIC metric is the maximum likelihood estimator. After fitting the ARIMA model, we generate predictions for every 3 months, six months and one year time steps to compute the residual value. Further, ARIMA residuals are modeled with NNAR(p, k) model having a pre-defined box-cox transformation set to λ = 0 to ensure the forecast values to stay positive. The value of p and k obtained by training the network and this is indeed a data dependent approach. Further, we add both the linear and nonlinear forecasts to obtain the final forecast results.

ARIMA(4,1,0) was fitted to the Philippines data having AIC = 1156.4 and log-likelihood value as −573.2. Further, the model residuals were trained using NNAR(18,10) model with an average of 20 networks, each of which is a 18-10-1 network with 201 weights. Then the forecast results of ARIMA along with NNAR residual forecasts are added together to obtain the final forecast values. And finally we compute RMSE, MAE, SMAPE and reported them in Table 2. For the San Juan dataset we fit an ARIMA(1,0,2) model achieving AIC = 8830.49 and log-likelihood=−4410.24 for the model. Further, an NNAR with p = 14 and k = 8 is trained. In a similar way as explained above, the final forecast values are computed and model performance metrics are reported in Table 3. Iquitos dataset fits an ARIMA(0,1,3) and its residuals are trained with NNAR(5,3) model. The quantitative measures of the proposed hybrid model and other state-of-the-art models are reported in Table 4. The estimated values of the proposed model for three datasets along with actual test values are depicted in Figure 2. It is clear from Table 2, 3, and 4 that the proposed hybrid ARIMA-NNAR model either outperforms single and hybrid models or remain competitive for three dengue datasets.

**Figure 2:**
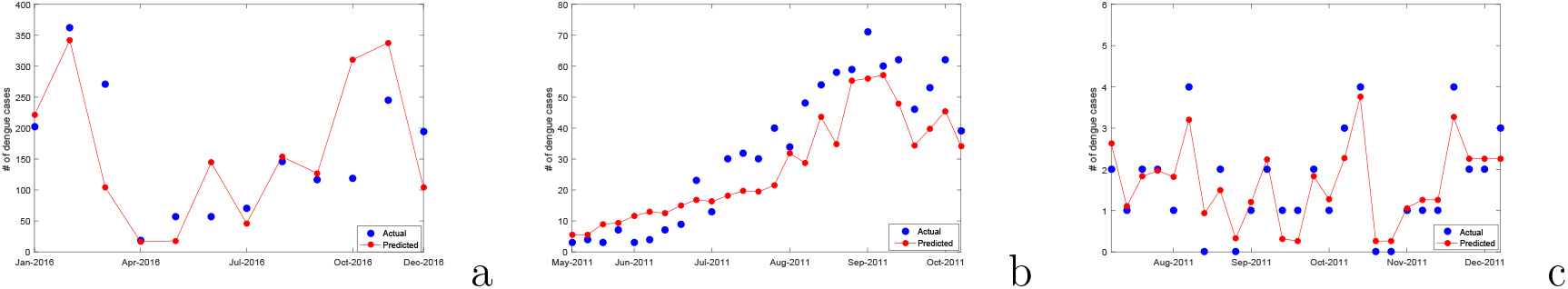
Actual vs predicted forecasts (using ARIMA-NNAR model) of the Philippines (a), San Jaun (b) and Iquitos (c) datasets

**Table 2:** Quantitative measures of performance for different forecasting models on the Philippines dataset

| Model | 6-Months ahead forecast |  |  | 1-Year ahead forecast |  |  |
| --- | --- | --- | --- | --- | --- | --- |
|  | RMSE | MAE | SMAPE | RMSE | MAE | SMAPE |
| ARIMA | 102.42 | 81.47 | 0.898 | 136.4 | 127.2 | 0.961 |
| ANN | 88.75 | 68.38 | 0.670 | 107.4 | 98.19 | 0.688 |
| NNAR | **57.28** | **47.24** | **0.441** | 97.57 | 75.55 | 0.689 |
| Hybrid ARIMA-ANN | 81.38 | 69.43 | 0.874 | 92.98 | 77.33 | **0.686** |
| Hybrid ARIMA-NNAR | 65.39 | 60.44 | 0.553 | **89.87** | **68.39** | **0.686** |

**Table 3:** Quantitative measures of performance for different forecasting models on San Juan dataset

| Model | 3-Months ahead forecast |  |  | 6-Months ahead forecast |  |  |
| --- | --- | --- | --- | --- | --- | --- |
|  | RMSE | MAE | SMAPE | RMSE | MAE | SMAPE |
| ARIMA | 7.801 | 7.230 | 0.636 | 21.68 | 17.59 | 0.668 |
| ANN | 9.511 | 6.991 | 0.577 | 26.33 | 20.79 | 0.765 |
| NNAR | 7.635 | 6.708 | 0.581 | 24.49 | 19.25 | 0.696 |
| Hybrid ARIMA-ANN | 7.781 | 7.238 | 0.635 | 21.48 | 17.45 | 0.663 |
| Hybrid ARIMA-NNAR | **7.438** | **6.569** | **0.570** | **20.73** | **16.56** | **0.612** |

**Table 4:**
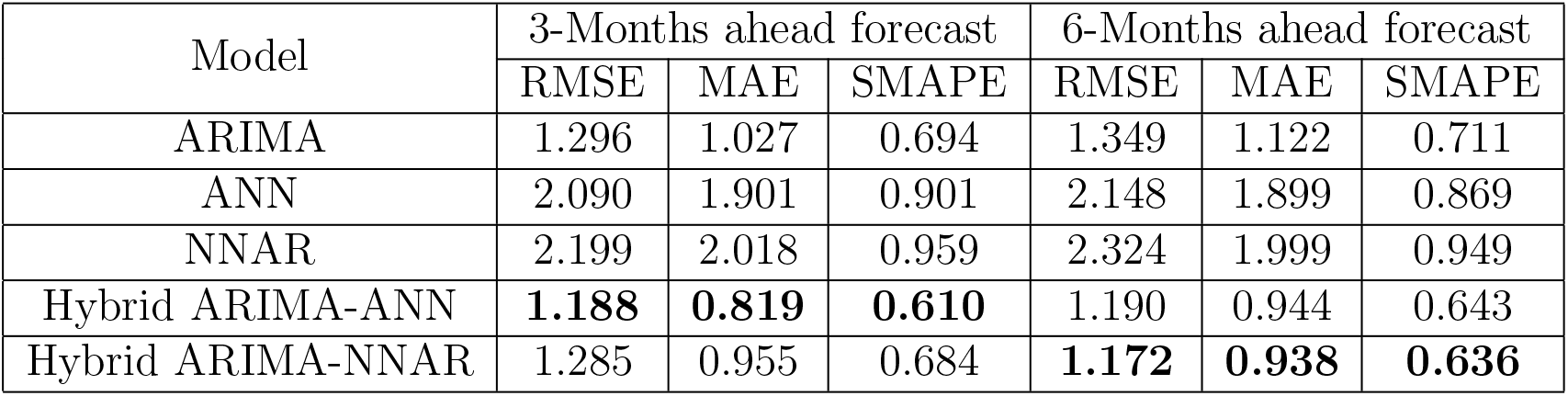
Quantitative measures of performance for different forecasting models on Iquitos dataset

| Model | 3-Months ahead forecast |  |  | 6-Months ahead forecast |  |  |
| --- | --- | --- | --- | --- | --- | --- |
|  | RMSE | MAE | SMAPE | RMSE | MAE | SMAPE |
| ARIMA | 1.296 | 1.027 | 0.694 | 1.349 | 1.122 | 0.711 |
| ANN | 2.090 | 1.901 | 0.901 | 2.148 | 1.899 | 0.869 |
| NNAR | 2.199 | 2.018 | 0.959 | 2.324 | 1.999 | 0.949 |
| Hybrid ARIMA-ANN | **1.188** | **0.819** | **0.610** | 1.190 | 0.944 | 0.643 |
| Hybrid ARIMA-NNAR | 1.285 | 0.955 | 0.684 | **1.172** | **0.938** | **0.636** |

## 4. Conclusion and Discussion

ARIMA is a well known classical time series model where as neural networks outperforms many of the nonlinear machine learning models in practice (29). The purpose of our empirical study was to build a model that performs superior for forecasting dengue epidemics. We proposed a hybrid ARIMA-NNAR model to filter out linearity using ARIMA model and predict nonlinear tendencies with the NNAR approach. In the development of the hybrid methodology, we need the component model to be sub-optimal and it will be effective to combine individual forecasts based on different information sets to produce superior forecasts. Hybrid ARIMA-NNAR model not only explains the linear and nonlinear autocorrelation structures present in the data better than traditional component and hybrid models but also yields better forecast accuracy than them. Though the limitation of the proposed methodology lies in the assumption of additive relationship between linear and nonlinear components. In addition, there is no theoretical guarantee that the residuals of ARIMA will always consist of valid nonlinearity of the data. It is true that no model can be universally employed in all circumstances, and this is in relevance with “*no free lunch theorem*” (30). Based on our experience on dengue forecasting, our model is best suited in the situations where the dataset will be quite large, high correlation among time series and there must be enough nonlinearity and non-stationarity present in the datasets. Our proposed model can be further used for seasonal time series data as well with small changes embed in the proposal, can be considered as a future research work of this paper. Finally, we can conclude that our proposed model can help policy makers to predict the subsequent dengue outbreaks accurately and respond to epidemics in a more effective way. Thus, this will reduce the impact of the future outbreaks and will govern the employment of resources.

## Acknowledgement

The authors gratefully acknowledge the financial assistance received from Indian Statistical Institute, Visvesvaraya Ph.D. Scheme and University Grants Commission, awarded by the Govt. of India.

## References

[1] O. J. Brady, P. W. Gething, S. Bhatt, J. P. Messina, J. S. Brownstein, A. G. Hoen, C. L. Moyes, A. W. Farlow, T. W. Scott, S. I. Hay, Refining the global spatial limits of dengue virus transmission by evidence-based consensus, PLoS neglected tropical diseases 6 (8) (2012) e1760.

[2] J. P. Messina, O. J. Brady, T. W. Scott, C. Zou, D. M. Pigott, K. A. Duda, S. Bhatt, L. Katzelnick, R. E. Howes, K. E. Battle, et al., Global spread of dengue virus types: mapping the 70 year history, Trends in microbiology 22 (3) (2014) 138–146.

[3] S. Bhatt, P. W. Gething, O. J. Brady, J. P. Messina, A. W. Farlow, C. L. Moyes, J. M. Drake, J. S. Brownstein, A. G. Hoen, O. Sankoh, et al., The global distribution and burden of dengue, Nature 496 (7446) (2013) 504.

[4] K. S. Vannice, A. Wilder-Smith, A. D. Barrett, K. Carrijo, M. Cavaleri, A. de Silva, A. P. Durbin, T. Endy, E. Harris, B. L. Innis, et al., Clinical development and regulatory points for consideration for second-generation live attenuated dengue vaccines, Vaccine 36 (24) (2018) 3411–3417.

[5] J. V. Silva, T. R. Lopes, E. F. de Oliveira-Filho, R. A. Oliveira, R. Durães-Carvalho, L. H. Gil, Current status, challenges and perspectives in the development of vaccines against yellow fever, dengue, zika and chikungunya viruses, Acta tropica.

[6] S. A. Lauer, K. Sakrejda, E. L. Ray, L. T. Keegan, Q. Bi, P. Suangtho, S. Hinjoy, S. Iamsirithaworn, S. Suthachana, Y. Laosiritaworn, et al., Prospective forecasts of annual dengue hemorrhagic fever incidence in thailand, 2010–2014, Proceedings of the National Academy of Sciences (2018) 201714457.

[7] T. K. Yamana, S. Kandula, J. Shaman, Superensemble forecasts of dengue outbreaks, Journal of The Royal Society Interface 13 (123) (2016) 20160410.

[8] P. Guo, T. Liu, Q. Zhang, L. Wang, J. Xiao, Q. Zhang, G. Luo, Z. Li, J. He, Y. Zhang, et al., Developing a dengue forecast model using machine learning: A case study in china, PLoS neglected tropical diseases 11 (10) (2017) e0005973.

[9] A. L. Buczak, B. Baugher, L. J. Moniz, T. Bagley, S. M. Babin, E. Guven, Ensemble method for dengue prediction, PloS one 13 (1) (2018) e0189988.

[10] L. R. Johnson, R. B. Gramacy, J. Cohen, E. Mordecai, C. Murdock, J. Rohr, S. J. Ryan, A. M. Stewart-Ibarra, D. Weikel, et al., Phenomenological forecasting of disease incidence using heteroskedastic gaussian processes: A dengue case study, The Annals of Applied Statistics 12 (1) (2018) 27–66.

[11] M. Gharbi, P. Quenel, J. Gustave, S. Cassadou, G. La Ruche, L. Girdary, L. Marrama, Time series analysis of dengue incidence in guadeloupe, french west indies: forecasting models using climate variables as predictors, BMC infectious diseases 11 (1) (2011) 166.

[12] V. Racloz, R. Ramsey, S. Tong, W. Hu, Surveillance of dengue fever virus: a review of epidemiological models and early warning systems, PLoS neglected tropical diseases 6 (5) (2012) e1648.

[13] A. E. Tümer, A. Akkuş, Forecasting gross domestic product per capita using artificial neural networks with non-economical parameters, Physica A: Statistical Mechanics and its Applications 512 (2018) 468–473.

[14] Y. Chen, B. Yang, J. Dong, A. Abraham, Time-series forecasting using flexible neural tree model, Information sciences 174 (3–4) (2005) 219–235.

[15] B. Zhu, Y. Wei, Carbon price forecasting with a novel hybrid arima and least squares support vector machines methodology, Omega 41 (3) (2013) 517–524.

[16] B. Kordanuli, L. Barjaktarović, L. Jeremić, M. Alizamir, Appraisal of artificial neural network for forecasting of economic parameters, Physica A: Statistical Mechanics and its Applications 465 (2017) 515–519.

[17] S. Arora, J. W. Taylor, Forecasting electricity smart meter data using conditional kernel density estimation, Omega 59 (2016) 47–59.

[18] M. Khashei, M. Bijari, A novel hybridization of artificial neural networks and arima models for time series forecasting, Applied Soft Computing 11 (2) (2011) 2664–2675.

[19] P.-F. Pai, C.-S. Lin, A hybrid arima and support vector machines model in stock price forecasting, Omega 33 (6) (2005) 497–505.

[20] L. Milacić, S. Jović, T. Vujović, J. Miljković, Application of artificial neural network with extreme learning machine for economic growth estimation, Physica A: Statistical Mechanics and its Applications 465 (2017) 285–288.

[21] J. Bozsik, Decision tree combined with neural networks for financial forecast, Periodica Polytechnica Electrical Engineering 55 (3–4) (2013) 95–101.

[22] G. Zhang, B. E. Patuwo, M. Y. Hu, Forecasting with artificial neural networks:: The state of the art, International journal of forecasting 14 (1) (1998) 35–62.

[23] R. J. Hyndman, G. Athanasopoulos, Forecasting: principles and practice, OTexts, 2018.

[24] M. R. Oliveira, L. Torgo, Ensembles for time series forecasting, Journal of Machine Learning Research.

[25] L. I. Kuncheva, Combining pattern classifiers: methods and algorithms, John Wiley & Sons, 2004.

[26] W. L. Gorr, Research prospective on neural network forecasting, International Journal of Forecasting 10 (1) (1994) 1–4.

[27] G. E. Box, G. M. Jenkins, G. C. Reinsel, G. M. Ljung, Time series analysis: forecasting and control, John Wiley & Sons, 2015.

[28] C. W. Granger, Combining forecaststwenty years later, Journal of Forecasting 8 (3) (1989) 167–173.

[29] N. K. Ahmed, A. F. Atiya, N. E. Gayar, H. El-Shishiny, An empirical comparison of machine learning models for time series forecasting, Econometric Reviews 29 (5–6) (2010) 594–621.

[30] D. H. Wolpert, W. G. Macready, No free lunch theorems for optimization, IEEE transactions on evolutionary computation 1 (1) (1997) 67–82.

